# High-throughput functional analysis of natural variants in yeast

**DOI:** 10.1101/2021.02.26.433108

**Authors:** Chiann-Ling C. Yeh, Andreas Tsouris, Joseph Schacherer, Maitreya J. Dunham

## Abstract

How natural variation affects phenotype is difficult to determine given our incomplete ability to deduce the functional impact of the polymorphisms detected in a population. Although current computational and experimental tools can predict and measure allele function, there has previously been no assay that does so in a high-throughput manner while also representing haplotypes derived from wild populations. Here, we present such an assay that measures the fitness of hundreds of natural alleles of a given gene without site-directed mutagenesis or DNA synthesis. With a large collection of diverse *Saccharomyces cerevisiae* natural isolates, we piloted this technique using the gene *SUL1*, which encodes a high-affinity sulfate permease that, at increased copy number, can improve the fitness of cells grown in sulfate-limited media. We cloned and barcoded all alleles from a collection of over 1000 natural isolates *en masse* and matched barcodes with their respective variants using PacBio long-read sequencing and a novel error-correction algorithm. We then transformed the reference S288C strain with this library and used barcode sequencing to track growth ability in sulfate limitation of lineages carrying each allele. We show that this approach allows us to measure the fitness conferred by each allele and stratify functional and nonfunctional alleles. Additionally, we pinpoint which polymorphisms in both coding and noncoding regions are detrimental to fitness or are of small effect and result in intermediate phenotypes. Integrating these results with a phylogenetic tree, we observe how often loss-of-function occurs and whether or not there is an evolutionary pattern to our observable phenotypic results. This approach is easily applicable to other genes. Our results complement classic genotype-phenotype mapping strategies and demonstrate a high-throughput approach for understanding the effects of polymorphisms across an entire species which can greatly propel future investigations into quantitative traits.

## Background

Quantitative traits, or traits that vary on a continuous distribution rather than in discrete categories, are responsible for most phenotypic differences across all organisms (MacKay et al., 2009; Morgante et al., 2018). Despite decades of efforts investigating how genotype informs phenotype, the molecular underpinnings of quantitative traits are still largely unknown, especially on a species-wide scale. Understanding how sequence changes lead to phenotypic changes is difficult to disentangle as these traits often involve interactions between multiple loci that each in themselves have genetic variation among natural populations. Even for a single locus, an exhaustive population-scale determination of genetic variants impacting phenotype remains out of sight. Improved approaches for investigating this in a cost-effective and high-throughput manner will greatly broaden insights into the genetic basis of trait variation, ranging from deleterious diseases to adaptive evolution.

Due to the rapid advancement and decreased cost of high-throughput sequencing, forward genetics approaches have boosted our ability to pinpoint loci underlying traits of interest. For instance, quantitative trait loci (QTL) and linkage mapping provide avenues for identifying loci responsible for phenotypic differences between individuals, and even in some instances can result in determining what polymorphisms are integral for certain phenotypes (Ehrenreich et al., 2012; Treusch et al., 2015). However, this method relies on pairwise crosses between a small subset of genetic backgrounds and is difficult to scale to investigate phenotypic variation on the population level (Stinchcombe and Hoekstra, 2008). Approaches like genome-wide association studies do investigate loci on the population scale, but lack the ability to infer and experimentally functionalize the effects of rare or low-frequency variants (Morgante et al., 2018). Other computational approaches deduce function based on collected data describing metrics such as conservation, statistical analyses of genomic architecture, allele frequency, predicted changes in protein stability, known sites of protein-protein interactions, and transcription factor-binding motifs, but all still require experimental validations (Adzhubei et al., 2010; Mitchell-Olds et al., 2007; Schymkowitz et al., 2005; She and Jarosz, 2018; Wagih et al., 2018; Wray et al., 2013).

Recently, multiplexed assays of variant effects (MAVE) studies have provided an approach for functionalizing thousands of variants in a high-throughput manner (Starita et al., 2017; Weile and Roth, 2018). These have been extremely useful in understanding how missense mutations or nucleotide changes alter gene function and/or expression (Duveau et al., 2017; Fowler and Fields, 2014; Matreyek et al., 2018). However, most of these approaches have been limited to studying single nucleotide or amino acid substitutions away from a reference sequence, as technologies don’t yet exist to generate and measure the consequences of the large libraries that would be necessary to explore combinatorial variation. Additionally, the majority of variants assayed are rarely reflective of those in natural populations. For instance, natural alleles can have more than one polymorphism, not all of which are seen exclusively in coding or exclusively in noncoding regions, and thus are not surveyed completely in many MAVE studies. Being able to directly test the function of natural variants of whole populations provides context for how polymorphisms and combinations of polymorphisms alter phenotype. Furthermore, such an approach would provide deeper insight into the evolutionary history of a gene and how both weak and strong selection or drift have acted upon a phenotype that results in the variation present in natural populations (Johnson and Barton, 2005; Mitchell-Olds et al., 2007). Thus, developing a method for testing natural variants in a high-throughput manner is of high interest.

Here, we developed such an assay functionalizing natural variants on a species-wide scale using *Saccharomyces cerevisiae*, the budding yeast. With the rapid advancement of high-throughput whole-genome sequencing, we now have large collections of natural *S. cerevisiae* strains that contain genomic data as well as geographical and ecological information (Bergström et al., 2014; Liti et al., 2009; Peter et al., 2018; Schacherer et al., 2009; Strope et al., 2015; Zhu et al., 2016). Although much research has been done on laboratory strains for understanding biology, curation of these collections revealed the striking diversity within this popular model organism: *S. cerevisiae* has been isolated from a variety of countries all over the globe and from habitats like human clinical samples, domesticated products like beer and bread, and tree and fruit samples. Sequencing of these genomes has revealed a lot about genetic variation, but still little is known about how these genetic changes impact phenotypic variation outside of a handful of association studies and QTL mapping efforts (Ehrenreich et al., 2012, 2009; Kim et al., 2012; Peltier et al., 2019; Wilkening et al., 2014). With the large genome sequencing efforts and strain collections, in addition to the wealth of molecular tools developed for yeast, *S. cerevisiae* is the ideal system to develop this assay and investigate the effects of natural polymorphisms for whole populations.

For piloting and developing our approach, we used the natural alleles of *SUL1* from a collection of 1,011 isolates to test whether we can deconvolute how variation affects cell growth under sulfate limiting conditions. *SUL1* encodes a high-affinity sulfate permease and is expressed under sulfate limitation. Previous studies have found that when evolving different strains of *S. cerevisiae* under sulfate limitation in the chemostat, cells with amplifications of *SUL1* have high fitness and rise in frequency in the population (Gresham et al., 2008; Payen et al., 2014; Sanchez et al., 2017). Strong selection for amplification of this locus in sulfate-limiting conditions allows for a reliable functional assay in which we can mimic amplifications by transforming cells with a low-copy plasmid containing *SUL1*. Additionally, we have previously performed a deep mutational scan on the promoter of *SUL1*, giving us a dataset measuring the functional consequences of single mutations for comparison (Rich et al., 2016). By co-culturing a population of cells transformed with a barcoded library of natural alleles, we can measure competitive fitness via barcode sequencing and thereby determine *SUL1* functionality *en masse*. Our results show that this assay is accurate in predicting function and useful in understanding what genetic changes affect phenotype. These data allow for insight into the evolutionary history of *SUL1* function and possible evidence for selection of loss-of-function mutations. This approach, especially when combined with established forward genetics approaches in identifying causal loci, will greatly strengthen our understanding of quantitative traits on a species-wide scale.

## Methods

### Strains and plasmids

Natural isolates from the 1,011 *Saccharomyces cerevisiae* collection were used to isolate natural variants of *SUL1* (Peter et al., 2018). Strains pinned on yeast extract peptone dextrose (YPD) agar plates were transferred to liquid YPD in 96-well plates, grown overnight at 30°C, and stored in 30% glycerol at −80°C. The FY3 S288C strain DBY7284 (*MATa ura3-52*) was used for transformation and competition experiments (described below). A GFP-marked strain YMD1214 (*MATa hoΔ::GFP-KANMX*) that has neutral fitness under sulfate limitation was used for validation competition assays. Prototrophic FY3 (DBY11069), YMD4321 (*MATa ura3-52 sul1Δ::URA3-KANMX*), YMD4322 (*MATa ura3-52 sul2Δ::URA3-KANMX*), and YMD4323 (*MATa ura3-52 sul1Δ::URA3-KanMX sul2Δ::URA3-KanMX*) were used to validate growth rates on sulfate-limited and sulfate-abundant agar plates. A pRS316 vector with an NruI site inserted in the BamHI site (YMD2307) was used in this study for molecular cloning and competitions described below. A complete list of strains can be found in **Supplementary Table 1**.

### Plasmid and yeast library generation

Strains from the 1,011 *S. cerevisiae* collection were pooled together from colonies on a solid agar plate. Genomic DNA was then extracted using the QIAGEN Genomic-tip 100/G kit. Natural variants of *SUL1* were amplified with primers designed to hybridize to conserved regions 844 bp upstream of the translation start site and 262 bp downstream of the stop codon (oligos 1 and 2, **(Supplementary Table 2**)). Oligo 2 also contained an 8 bp randomized sequence to serve as a barcode. PCR was performed using KAPA HiFi Hotstart Readymix with the following cycling conditions: 95°C for 3 min, then 19 cycles of 98°C for 20 seconds, 60°C for 15 seconds, and 72°C for 4 minutes. Final extension was at 72°C for 4 minutes, and then the reaction was cooled to 4°C. The barcoded product was purified using the DNA Clean and Concentrator kit from Zymo Research and assembled into an NruI-digested plasmid via Gibson assembly. Chemically competent *E. coli* cells were transformed with the product using heat shock at 42°C, and >20,000 transformants were collected and pooled. Plasmids were extracted from the pooled transformants using Wizard® *Plus SV* Miniprep DNA Purification Kit and then used to transform yeast (DBY7284) using 100 μL of 2 M lithium acetate, 800 μL of 50% 4000 polyethylene glycol, 100 μL of 1M dithiothreitol, and 50 μL of 10 mg/mL of carrier DNA. Approximately 6,000 Ura+ yeast transformants were collected for pooled competition experiments and for PacBio sequencing (**Figure 1A**).

**Figure 1.**
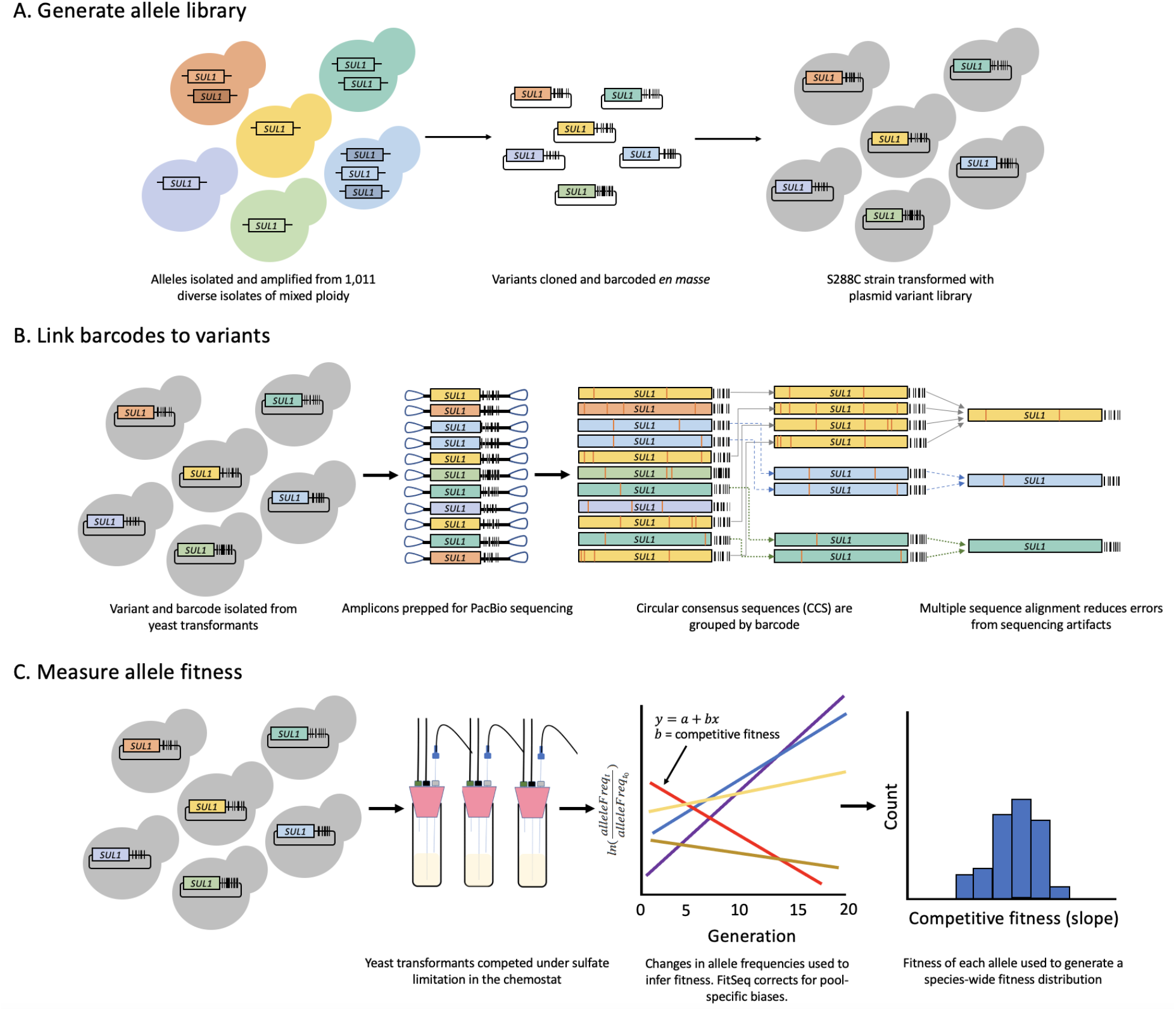
Workflow for assaying natural variants in the 1,011 strain collection. **A)** The S288C lab strain is transformed with *SUL1* natural allele barcoded plasmid library. **B)** PacBio long-read sequencing is used to link barcodes with variants. **C)** Transformants are competed together under sulfate limitation. Barcode sequencing every 3-4 generations is used to calculate the abundance of each variant and its respective competitive fitness.

### Linking barcodes with full-length variants

Plasmids were extracted from the yeast transformant pool using Zymoprep Yeast Plasmid Miniprep II (Zymo Research). Plasmid fragments containing the barcode and variant were isolated using M13/pUC primers with KAPA HiFi Hotstart Readymix and the following cycling conditions: 95°C for 3 min, then 13 cycles of 98°C for 20 seconds, 60°C for 15 seconds, and 72°C for 4 minutes. The final product was extracted from a 0.5% agar gel using Qiagen’s Gel Extraction kit and cleaned using Ampure PB beads (Pacific Biosciences). Two PacBio libraries were made using the SMRTbell™ Template Prep Kit 1.0 (Pacific Biosciences) and sent to University of Washington PacBio Sequencing Services for sequencing and Sequel II circular consensus sequence (CCS) analysis.

BAM files of CCS reads were aligned to the plasmid reference file using BWA/0.7.13 mem (Li, 2013). Reads that were aligned to the reference were piped to a new BAM file with Samtools/1.9 (Li et al., 2009). These reads were also analyzed with cigar strings to validate alignment of PacBio reads. From there, the barcodes were extracted, and a barcode-variant map was generated that contained a file with all of the barcode-variant reads and all of the highest quality reads for each barcode, as previously described (Matreyek et al., 2018). Since the resulting barcode-variant map still showed a considerable number of insertion and deletion errors, we used a multiple sequence alignment of all the reads that shared the same barcodes to eliminate additional sequencing errors. Alignments were done using MUSCLE (v.3.8.31) (Edgar, 2004). Any further ambiguous nucleotides were resolved by performing a pairwise alignment against the highest quality read (EMBOSS Needle v. 6.4.0) (Needleman and Wunsch, 1970).

To match PacBio reads to strains in the 1,011 collection, reference sequences were first extracted from the GVCF in the 1,011 collection genome data using BCFtools consensus. We then used regular expressions to search for reads that were putatively derived from these reference sequences. We removed barcodes that contained only one CCS read or were not represented in our barcode sequencing analysis (**Figure 1B**).

### Pooled library competition in chemostats

Sulfate-limited media (3mg/L ammonium sulfate) was prepared as previously described (Gresham et al., 2008; Payen et al., 2014). Four 50 mL chemostat culture vessels were filled with 20 mL of media at 30°C and inoculated with 1 mL of the yeast transformant pool. This culture was grown for 24 hours, after which the pumps were turned on and the culture switched to a continuous culture system at a dilution rate of about 0.17 volumes per hour (∼3.4 mL/hour in a 20 mL culture). Samples were taken twice a day for 5 days, or about 25 generations. For each sample, 1 mL was stored in 25% glycerol at −80°C, and another 1 mL was used for plasmid extraction (**Figure 1C**).

### Barcode sequencing and analysis

For each time point and replicate from the pooled library competition, plasmids were again extracted using the Zymoprep Yeast Plasmid Miniprep kit. One replicate was discarded due to technical errors. Barcodes were isolated and amplified using forward oligo 25 and indexed reverse oligos 26-40 and 120-128 that included Illumina Nextera sequencing adaptors. KAPA HiFi Hotstart Readymix was used with 1 μL of 1X SYBR™ Green I and the following PCR cycles: 95°C for 3 min, then 17-19 cycles of 98°C for 20 seconds, 60°C for 15 seconds, and 72°C for 15 seconds. The reaction was run on a Bio-Rad MiniOpticon (Bio-Rad) to avoid overamplification. PCR products were cleaned using Ampure XP Beads (Agencourt) and quantified using the KAPA Library Quantification Kit for Illumina® Platforms (Roche). Libraries were sequenced on a NextSeq sequencer (Illumina) with sequencing oligos 41 (Read 1), 44 (Read 2), and 100 (Index). Paired-end reads were merged using PEAR/0.9.5 (Zhang et al., 2014). Using FitSeq, we calculated the fitness of each barcode for each given replicate (Li et al., 2018). FitSeq normalizes each pool to account for experimental error between replicates, providing a more accurate readout of fitness. The fitness values were then normalized by the average fitness of barcodes associated with the wild-type (S288C) alleles. The effects of single mutations were also compared with predicted consequences of mutations from mutfunc (Wagih et al., 2018) (**Figure 1C**).

### Pairwise fitness assays in chemostats

To assess fitness of strains carrying individual alleles, 300 μL of a liquid culture of each strain was inoculated into a chemostat containing 20 mL of sulfate-limited media at 30°C. For each of the competitors, one competitor strain contained a plasmid with an extra copy of *SUL1* and the other isogenic strain contained a neutral GFP marker. Additionally, competition experiments of strains carrying each allele being assayed were conducted in at least two biological replicates. Cultures were grown for 24 hours before switching to a continuous culture system. Once cultures achieved steady state, the competing cultures were mixed at a 1:1 ratio. Cultures were competed for 15 generations after mixing and sampled twice daily (approximately every 3-6 generations). For each sample, cultures were assayed for percent GFP cells with a BD Accuri C6 flow cytometer (BD Biosciences). Competitive fitness values were calculated by plotting ln(number of dark cells/number of GFP+ cells) over about 25 generations and taking the linear slope of the linear regression from this data.

### Measuring the growth of the 1,011 isolates on solid media

Solid sulfate-limited media (3mg/L ammonium sulfate) was prepared by adding 2% agarose to liquid sulfate-limited media and poured in PlusPlates (Singer Instruments). Solid sulfate abundant media was prepared by adding ammonium sulfate (to 5g/L) to the sulfate-limited media. To ensure the depletion of sulfate in the cells, all 1,011 natural isolates were grown overnight (∼14 hours) on solid sulfate-limited media. The isolates were then replicated in quadruplicate on solid sulfate-limited and sulfate-abundant media. Photos of the colonies were taken every 12 hours for 3 days and the R package gitter was used to calculate the size of each colony in the photos (Wagih and Parts, 2014). For each time point on both limited and abundant conditions, we subtracted the colony size at the first time point from the colony size at subsequent time points (colony size = size_t_ - size_t=0_). Growth rates were calculated by taking the average of the ratio of the colony size in limited media over the colony size in abundant media across 72 hours.

### Phylogenetic tree generation and sequence analysis

To generate our phylogenetic trees, *SUL1* sequences from the 1,011 strains and from *S. paradoxus* strain CBS432 were aligned using MUSCLE (Edgar, 2004). The genetic distances for *SUL1* alleles were calculated using the maximum-likelihood-based distances through DNADIST in the PHYLIP package (Felsenstein, 2005). A gene tree for *SUL1* was then generated using the NEIGHBOR program, and the final tree was visualized and annotated using R/ggtree (Yu et al., 2017).

To determine the prevalence of loss-of-function mutations across all 1,011 strains, we used sequences from the core ORFs in the pangenome as reference sequences and identified which strains were homozygous for premature stop codons in each of the core ORFs (Peter et al., 2018). Premature stop codons that occurred in the last 90% of an ORF were not included, as previous studies have shown that these mutations would not necessarily cause a significant loss of function (Bergström et al., 2014). Gene Ontology (GO) analysis was conducted using Yeastmine (accessed May 22, 2020) and both Benjamini-Hochberg and Bonferroni test corrections were used to account for multiple testing (Balakrishnan et al., 2012).

### Data availability

Raw sequencing data can be found in the Sequencing Read Archive (BioProject Accession PRJNA681436 https://www.ncbi.nlm.nih.gov/bioproject/PRJNA681436). Scripts and Supplemental Tables used for this paper can be found at https://github.com/dunhamlab/SUL1_natural_variants. All alleles, matched strains, barcodes, fitness, coding mutations, and noncoding mutations can be found in **Supplementary Table 3**.

## Results

### Allele library curation and characterization

When sulfate is a limiting nutrient, *S. cerevisiae* increases expression of *SUL1*, which encodes a high-affinity sulfate permease that increases the intake of sulfate molecules into the cell. Previously, we measured the competitive fitness of *SUL1* alleles isolated from 10 different wild yeast isolates (Payen et al., in preparation). We found that these alleles confer a wide range of fitness: some had loss-of-function phenotypes while others performed better than the allele found in the reference strain S288C. In order to determine if this wide variation was representative across the entire species and whether it correlated with features such as the environment from which each strain was isolated, we set out to survey *SUL1* functionality across a bigger sample of natural isolates. For this study, we used the 1,011 *S. cerevisiae* strain collection, which was curated from a variety of geographical and ecological origins (Peter et al., 2018). In addition, the collection contains at least 250 unique alleles of *SUL1* with 354 variable sites in the gene. Alleles contain 11 polymorphisms on average vs. the reference allele, with the most polymorphic allele having 79 mutations. Therefore, these factors make *SUL1* a powerful tool for us to better understand the natural variation of a single gene in *S. cerevisiae* populations.

In our previous studies, the fitness of individual *SUL1* alleles was measured by transforming the reference strain with an additional copy of a *SUL1* allele on a low-copy plasmid, and the resulting strain was competed against an isogenic GFP-marked strain under sulfate limitation (Payen et al., in preparation) (Sanchez et al., 2017). While this assay is reliable and consistent, it would be difficult and unrealistic to scale to measure hundreds of alleles. Thus, we developed a high-throughput, multiplexed approach that allows us to simultaneously measure these fitness values directly (**Figure 1**). To do this, we pooled the 1,011 isolates together, extracted genomic DNA, and used barcoded primers binding to conserved regions to isolate and amplify all natural alleles of the *SUL1* gene. These sequences were cloned *en masse* onto low-copy CEN/ARS plasmids to create an allele library and used to transform the reference strain (FY). The resulting library contained approximately 6,000 barcodes for an estimated 250 unique alleles (24X coverage) to ensure complete coverage and internal replicates (**Figure 1A**).

We used PacBio long-read circular consensus sequencing (CCS) to pair barcodes with their respective alleles (**Figure 1B**). Although PacBio CCS has drastically improved and decreased sequencing errors over the past few years, we found that many reads still contained errors that were especially noticeable in the form of insertions and deletions. To further eliminate these sequencing artifacts, we performed multiple sequencing alignments on CCS reads that shared the same barcode, and used those to derive new consensus sequences. In total, our analysis produced 8,386 barcode-variant pairs, which we determined was still an overestimate given our library size of ∼6,000 barcodes. We removed consensus reads that only appeared once to eliminate false positives or negatives in our downstream analysis, with 3,787 barcodes remaining.

Among these 3,787 barcodes, we identified 407 unique alleles in our library. Of these variants, we were able to match 228 alleles to at least one strain in the 1,011 strain collection, with a total of 880 strains that had at least one matched allele in the library (**Supplemental Figure 1**). To determine how well this library reflected the polymorphisms in the strain collection, we plotted the correlation of polymorphism frequency in both the variant reference sequences and library sequences and found that these values were highly correlated (Pearson’s correlation, r=0.978, **Supplemental Figure 2**). Correlation values were similar for polymorphisms in all regions of the gene: the 5’-UTR, coding region, and 3’-UTR were all well-correlated (Pearson’s correlation, r=0.956, 0.980, and 0.993, respectively). Of the 354 variable sites found in the reference sequences, only 45 of them were not detected in the allele library, nine of which were rare polymorphisms. Our pipeline did not reveal any *de novo* mutations that could have resulted from PCR or sequencing artifacts.

We were also unable to capture the alleles from 23 strains that were identified to have *SUL1* introgressed from *Saccharomyces paradoxus*. This was likely due to these sequences being more highly diverged and therefore unable to hybridize with the primers that were designed. However, for completeness, we were still able to measure the functionality of the introgressed *SUL1* alleles using our lower throughput method of direct competitions, as described below.

### Fitness distribution across natural SUL1 alleles

To determine the fitness landscape of all the *SUL1* alleles present in our allele library, we competed the library of yeast transformants in a continuous culture system under sulfate-limited media. Samples from 12 timepoints across four replicates were collected every 3-4 generations. For each sample, we extracted the plasmids from sampled cultures and sequenced the ba rcodes using Illumina short-read sequencing. By tracking the change in barcode frequencies over the 12 timepoints, we determined the competitive fitness values for strains carrying each allele (**Figure 1C**). The calculated competitive fitness of the three replicates showed strong correlation and reproducibility (**Supplemental Figure 3**).

In our barcoded library, 863 of 3,787 barcodes were associated with alleles identical to that of the S288C reference strain. We normalized all fitness values to the average fitness of these wild-type alleles (0.0097, standard deviation 0.0698). Reassuringly, we found that many barcodes with lower fitness values (fitness < −0.03) were largely associated with alleles containing natural premature stop codons (**Figure 2**). In fact, upon analyzing the sequences in each strain, we found 74 strains that are homozygous for premature stop codons in their *SUL1* alleles. Among the 31 alleles with premature stop codons, fifteen occur in amino acid positions 155 and 184 (Y155* and 184Q*, where amino acids are compared to the S288C protein sequence).

Due to the wide range in fitness of alleles with premature stop codons, we investigated whether stop codons that occurred earlier in *SUL1* have a greater impact on function. We found that the location of stop codons in *SUL1* did not dictate the deleterious effects of containing a nonsense mutation (**Supplemental Figure 4a**). However, the fitness of alleles with premature stop codons at amino acid position 671 consistently have much lower fitness compared to others with premature stop codons elsewhere. This stop codon occurs in the predicted extracellular STAS (sulfate transporter and anti-sigma factor antagonist) domain, which is thought to be crucial for metabolism sensing, and may be further impacting sulfate transport under sulfate limiting conditions (Sharma et al., 2011).

We compared the standard deviations among barcodes that shared the same loss-of-function alleles to that of barcodes that shared the same wild-type alleles (**Supplemental Figure 4b**). The barcodes linked to loss-of-function alleles do vary more in fitness (Welch two sample t-test, p < 0.005), although we attribute this variance to increased errors that occur when measuring fitness on a log scale. In regard to magnitude, the barcode counts are reliable, but the barcode counts tend to be less accurate when frequencies are low and continue to decrease through later time points.

In addition to stratifying alleles with premature stop codons and alleles with wild-type phenotypes, we identified alleles with nonsynonymous polymorphisms that also result in a loss of function. For instance, alleles that have a single polymorphism resulting in a T669K amino acid substitution show a loss of function. We also found that alleles with A454P and D483N and alleles with S699L substitutions (and no additional nonsense or promoter polymorphisms) have a loss of function phenotype in our pooled library. Alleles with their polymorphism information, corresponding strain information, and measured fitness values can be found in **Supplementary Table 3**.

We assessed how well the fitness values are reflected in direct competitions by selecting *SUL1* alleles from seven isolates and cloning them individually on the same low-copy plasmid. We transformed S288C haploid yeast with these individual plasmids and competed each allele directly against an isogenic GFP strain with no plasmid (**Figure 3**). Three of the alleles were selected to validate a wild-type-like phenotype and corresponded to the values calculated in the pooled competition. Three other alleles selected showed a loss-of-function phenotype in the pooled competition, which was reflected in the direct competitions. Two of these alleles contained a deletion that resulted in a frameshift (from strains BGM and AQM), and the third allele had nonsynonymous mutations (from strain BII). The BII strain has previously been evolved through sulfate limitation for 150 generations, and it was found that a natural polymorphism that results in a P296L change is responsible for the loss-of-function phenotype (Payen et al., in preparation). In each case, we found the results of the direct competitions recapitulated those found in our pooled competition.

Since we were unable to measure functionality of introgressed alleles in our library, we used the same approach of a direct competition to assay introgressed allele functionality. After validating the fitness of the *SUL1* orthologue from *S. paradoxus* in the *S. cerevisiae* background, which has previously shown high fitness (Sanchez et al., 2017), we also tested the fitness of two alleles that show signatures of introgression from *S. paradoxus*. The two introgressed alleles, despite having over 40 amino acid differences compared to the reference allele, also have a wild-type phenotype (**Figure 3**).

**Figure 3.**
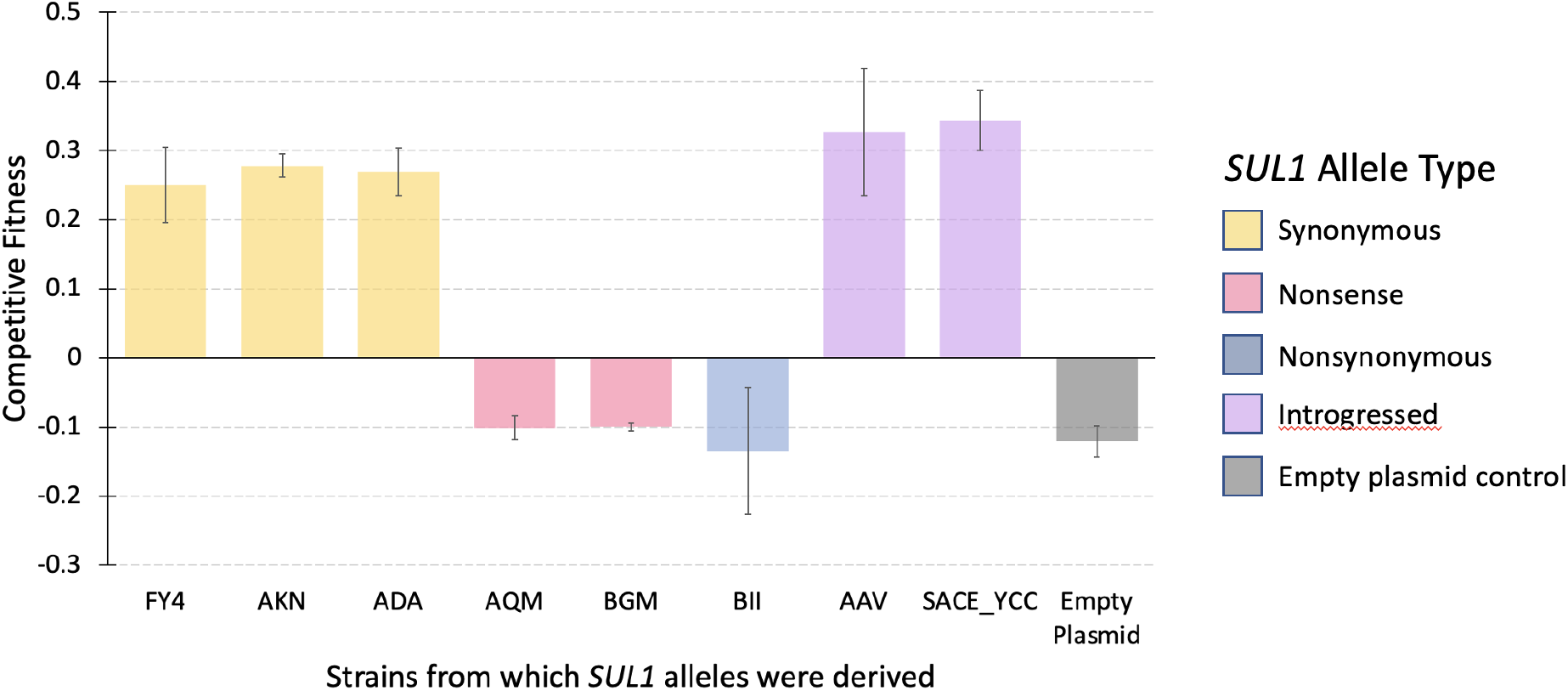
Validation of pooled competition through direct competitions of selected natural *SUL1* alleles. S288C strains transformed with a specific *SUL1* allele on a low-copy plasmid were individually competed against an isogenic GFP-marked strain in the chemostat. Fitness values were calculated by tracking the log-fold change in proportion of non-fluorescent strains and fluorescent strains over 20 generations. These values were used to validate select alleles and their phenotypes observed in the pooled competition. Alleles were selected based on definitive categorization in wild-type-like (pooled competitive fitness close to 0) or loss-of-function (pooled competitive fitness less than −0.10) phenotypes. Of the loss-of-function alleles, AQM and BGM have premature stop codons while BII is loss-of-function due to nonsynonymous polymorphisms. *SUL1* alleles in AKN, ADA, and BII were done in a prior experiment (Payen et al., in preparation).

### Effects of promoter mutations in natural SUL1 variants

The fitness distribution across the natural alleles shows alleles with only synonymous site changes in the coding region that nevertheless have a lower competitive fitness compared to strains carrying the wild-type coding sequence from the reference strain (**Figure 2**). We reasoned that these alleles may instead carry functional differences in the noncoding sequences. We found that these alleles share the n.-456G>A polymorphism, and upon further inspection discovered that this SNP is only present in alleles (including those with additional nonsynonymous SNPs) with lower competitive fitness values under sulfate limitation (median competitive fitness = −0.04). Since this competitive fitness value is not as low as alleles with premature stop codons (median competitive fitness = −0.17), it is indicative of an intermediate phenotype. This SNP occurs in a putative Cbf1-binding motif, and binding of this Cbf1 transcription factor has been shown to be important for growth in sulfate limiting conditions (Rich et al., 2016; Siggers et al., 2011). The SNP also decreased fitness in a *SUL1* promoter mutagenesis study, further supporting the functional effects of changes in this motif (Rich et al., 2016).

**Figure 2.**
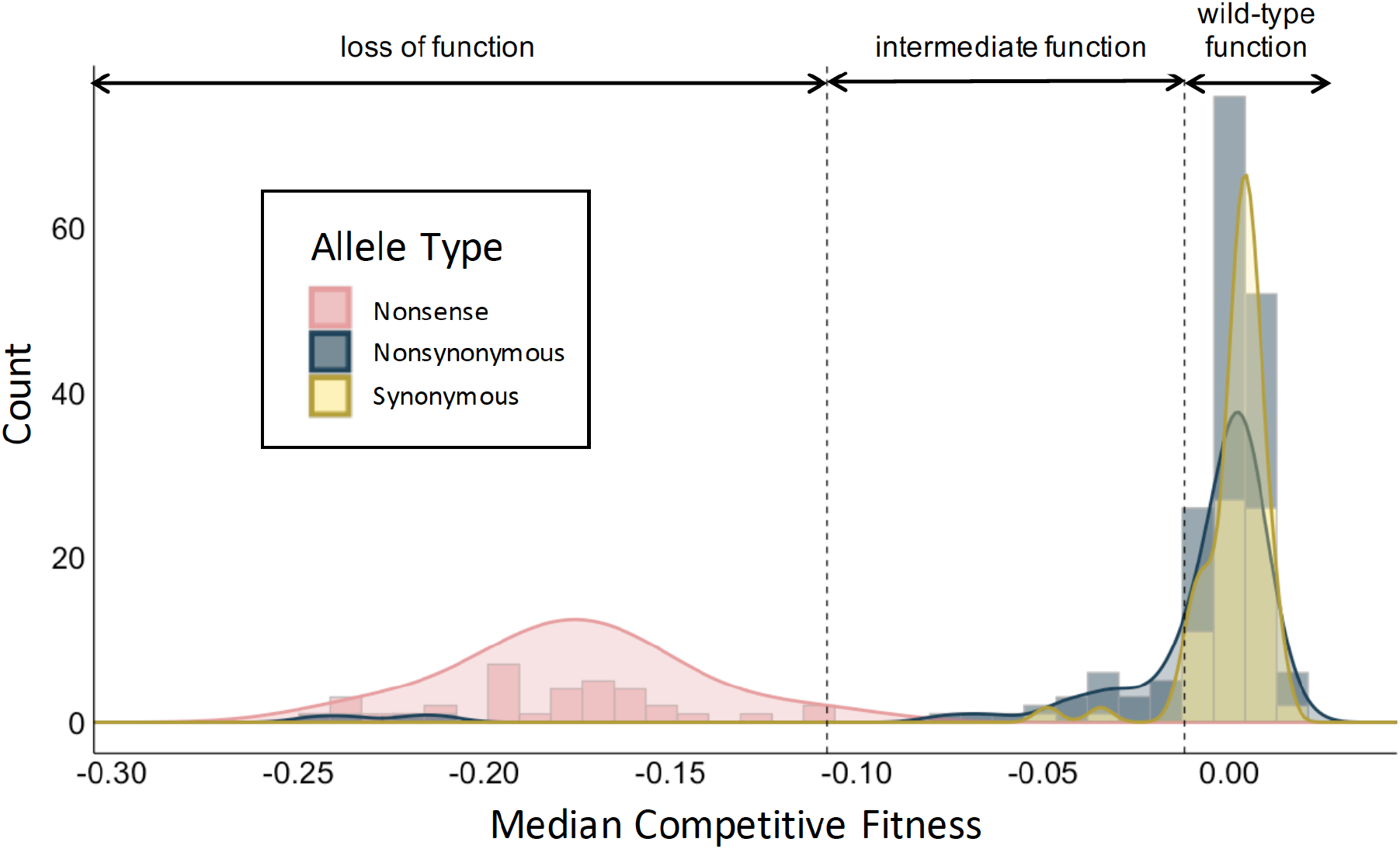
Species-level distribution of fitness effects of natural *SUL1* alleles. Lab strain S288C yeast transformed with an allele library of *SUL1* cloned onto a low-copy plasmid were competed in sulfate-limited media in the chemostat. The log-fold change in proportions of each barcode across 12 timepoints were measured through barcode sequencing and used to calculate competitive fitness. Alleles categorized as nonsense alleles may also contain synonymous and nonsynonymous polymorphisms. Those grouped as nonsynonymous alleles may contain synonymous polymorphisms, but do not have premature stop codons. Synonymous alleles do not have nonsynonymous or nonsense polymorphisms. All alleles may contain polymorphisms in the promoter or 3’UTR. Loss-of-function alleles were defined as having a fitness lower than the highest-fit allele with a premature stop codon. Wild-type function alleles have a fitness higher than the lowest-fit synonymous allele.

We used the highest fitness of an allele that contains a premature codon (median competitive fitness = −0.108) and the lowest fitness of alleles without promoter or nonsynonymous polymorphisms (median competitive fitness = −0.0120) to establish a range for other alleles with intermediate phenotypes. Twenty-two unique alleles show an intermediate phenotype, and 9/20 alleles with nonsynonymous polymorphisms also have the n.-456G>A polymorphism. Using these benchmarks, we also identify nonsynonymous changes that do not confer a complete loss of function.

The observation of promoter mutations affecting phenotype in sulfate limitation led us to inspect how much promoter polymorphisms in general contribute to the fitness values observed across the entire allele library. We compared the standard deviation in fitness for sequences that share the same coding sequence to the standard deviation in fitness for sequences that share the same promoter sequences. We found that the coding sequences seemed to more consistently determine fitness of a strain under sulfate limitation (**Figure 4a**). That is, alleles with the same promoter sequences had a greater variance in fitness values. Furthermore, alleles that shared the same coding sequences but differed in promoter sequences showed few significant differences in fitness (**Figure 4b**). Finally, despite the fact that the promoter mutagenesis study found mutations that could improve fitness under sulfate limitation, we did not identify such polymorphisms among our natural variants.

**Figure 4.**
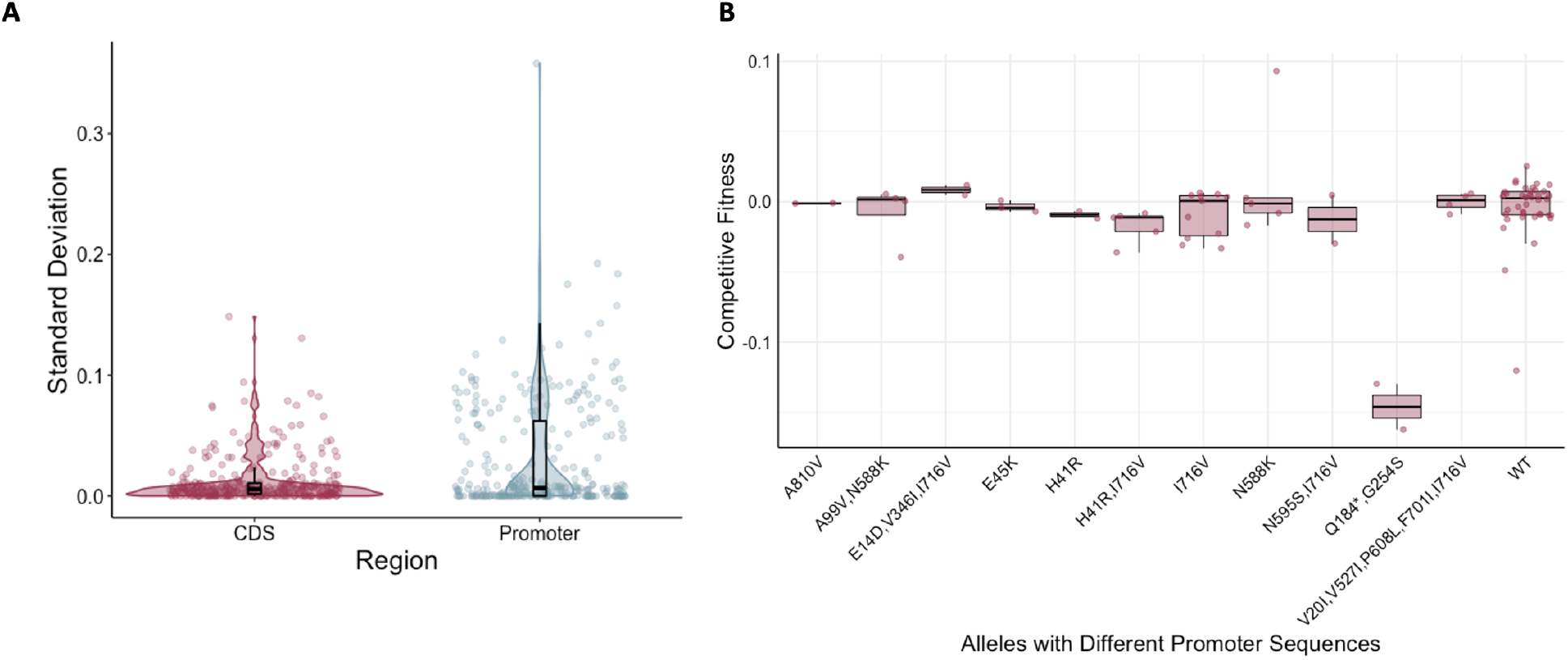
Coding polymorphisms are more useful for predicting deleterious effects compared to those in the promoter of *SUL1*. **A)** Violin plots of the standard deviations of the competitive fitness for barcodes grouped by those that share the same coding sequence compared with the standard deviation of those that share the same promoter sequence. **B)** Boxplots of competitive fitness of the sequences that share the same coding sequence but differ in the promoter sequences.

### Comparing competitive fitness with mutfunc

With nonsense mutations, loss of function can be predicted based on sequence alone. However, predicting the functional effects of other mutations based on sequence alone is much more challenging. To determine how well these fitness values were reflected in functional computational predictors, we used mutfunc to compare our results to predicted functional effects. For each variant, we took the most putatively detrimental mutation and compared its value to the fitness values calculated in our pooled competition assay. While the SIFT scores and our fitness values themselves showed very little correlation (**Supplementary Figure 5a**), we found that most alleles with a loss-of-function phenotype had a low SIFT score (**Supplementary Figure 5b**). Interestingly, many mutations that SIFT predicted would be detrimental actually had a wild-type-like phenotype under sulfate limitation. This highlights the value of experimentally measuring the function of variants, especially in cases where we need to consider the functional impacts of multiple polymorphisms on the same haplotype.

### Phylogenetics and sequence analysis of natural SUL1 alleles

To assess phenotypic patterns of *SUL1* on the population level, we annotated a distance-based gene tree of *SUL1* (**Figure 5**) with the competitive fitness values we calculated from our pooled competition assay. In our gene tree, we used the *SUL1* allele of *Saccharomyces paradoxus* (CBS432) as the outgroup. We removed branch lengths from these trees to simplify interpretations. Using these annotated trees, we are able to interpret phenotype in relation to ecological origins and phylogenetic relationships (**Figure 5**). We firstly looked at the strains homozygous for premature stop codons in *SUL1*. The polymorphism that results in Q184* does not occur in a singular clade, reducing the possibility that this premature stop codon arose in prevalence as a result of drift or identity by descent. Alleles with Y155* are primarily present in strains isolated from dairy environments in Normandy, France; however, not all dairy strains share the same nonsense mutation (**Figure 6**). Two other strains derived from dairy, AQM and BGM, instead have the L125* frameshift mutation. This pattern suggests that a loss-of-function mutation could be beneficial in a dairy environment.

**Figure 5.**
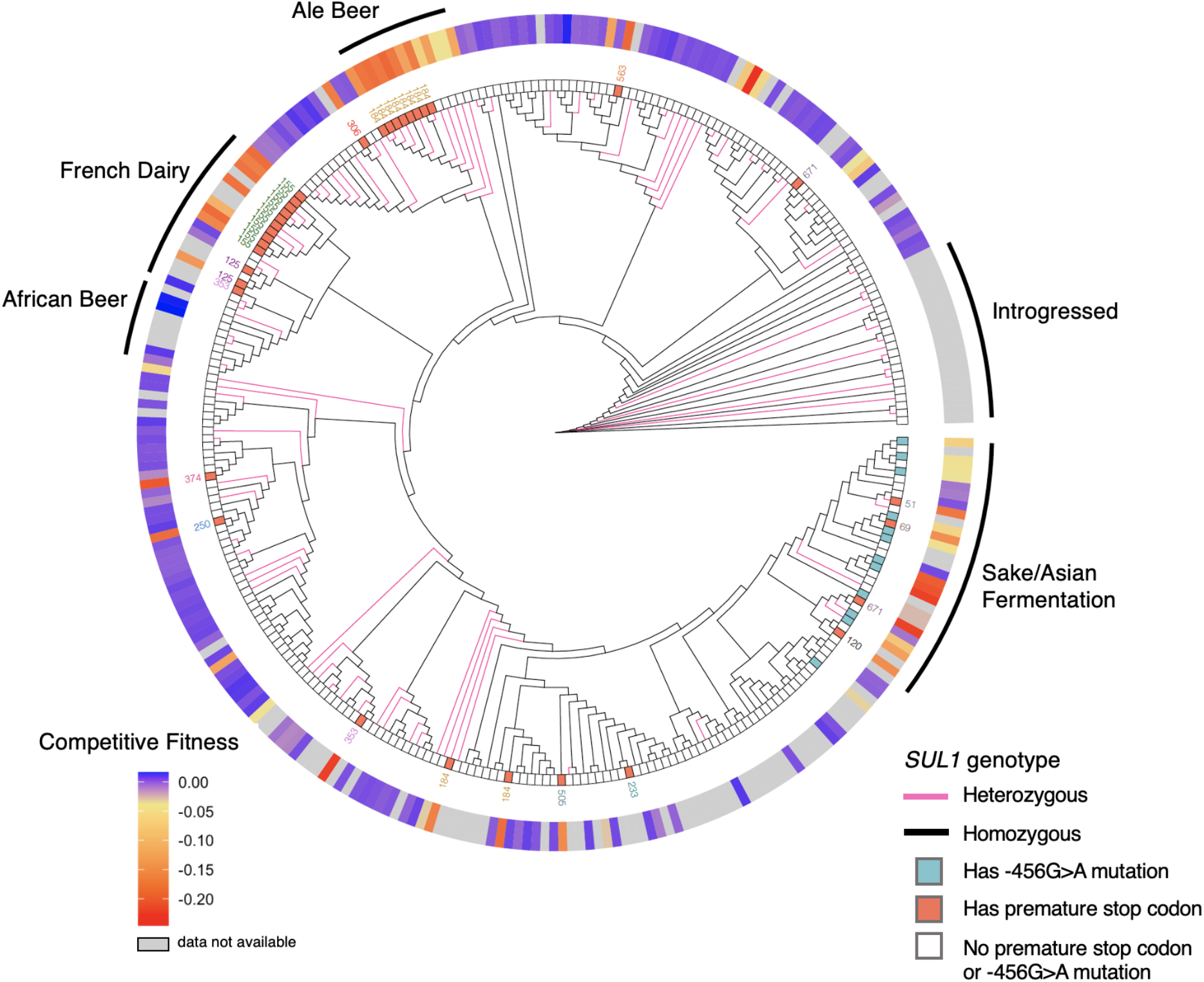
Neighbor-joining gene cladogram generated through PHYLIP using unique genotypes of *SUL1* in the 1,011 strain collection. French dairy and sake/Asian fermentation clades both show multiple independent instances of loss-of-function mutations. A stop codon at amino acid position 184 occurs independently in different strains. Color of edges (pink or black) indicates whether genotype for those terminal nodes are homozygous or heterozygous. Heterozygous alleles can be derived from diploid, triploid, tetraploid, or even pentaploid strains. Boxes directly adjacent to terminal nodes indicate the genotypes that are homozygous for a premature stop codon (red) or a −456G>A mutation (cyan). Flanking boxes of genotypes with premature stop codons are numbers indicating where in the amino acid sequence the premature stop codon occurred. The ring surrounding the tree denotes the mean *SUL1* competitive fitness values for a given strain’s allele on a purple (wild-type-like fitness) to red (loss-of-function fitness) gradient. Labeled regions are generalizations for what comprises most of those clades.

**Figure 6.**
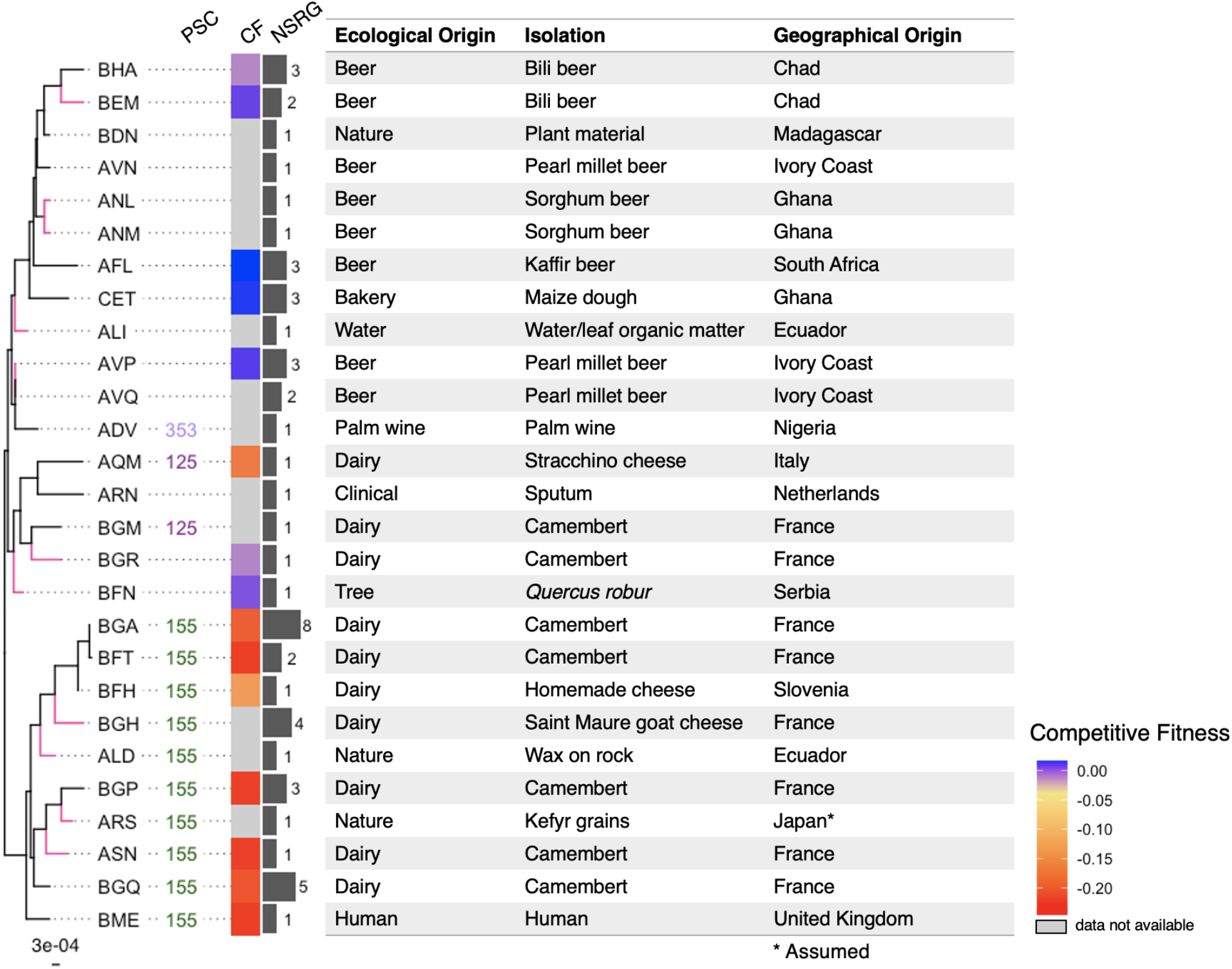
Dairy and African beer subtree of the 1,011 *SUL1* genotypes. Although dairy strains AQM and BGM share a more recent common ancestor to African beer strains, they show different but independent and homozygous loss-of-function polymorphisms. Color of edges (pink or black) indicate whether genotype for those terminal nodes are homozygous or heterozygous. PSC, amino acid site with premature stop codon (homozygous); CF, competitive fitness; NSRG, number of strains represented by genotype. Boxes around terminal nodes indicate the genotypes that are homozygous for a premature stop codon (red) or a −456G>A mutation (cyan). Scale (bottom left) indicates number of nucleotide substitutions per site.

The majority of strains with the detrimental promoter mutation n.-456G>A were isolated from sake or Asian fermentation strains. Additionally, many strains in this clade have a premature stop codon and or nonsynonymous polymorphisms that result in loss of function, which would again support the idea that there may be a trade-off for having a loss-of-function *SUL1* allele since more than one loss-of-function allele sequence exist among these strains.

Based on the distribution of deleterious alleles over the phylogeny, we wondered if these allele differences would lead to phenotype differences when the alleles were in their native strain context. We grew all isolates (unmodified) from the 1,011 strain collection on solid minimal media agar plates under sulfate limitation and compared the growth rates to that of the strains pinned on sulfate-abundant minimal media. Interestingly, we found little to no correlation between the growth rates of strains and the competitive fitness values of their *SUL1* alleles (**Supplemental Figure 6a**,**b**). We additionally looked for growth patterns among ploidy, geographical origins, and clade and found no patterns related to these groupings (**Supplemental Figure 6c**). These results argue that additional background effects beyond the *SUL1 locus* matter for determining fitness in sulfate limitation. Measuring the fitnesses of the allele library in additional strain backgrounds may help further characterize this genetic complexity.

We calculated the average *dN/dS* value of *SUL1* across all 1,011 strains and found that the value was low (*dN/dS* < 0.2), suggesting that there may be purifying selection on *SUL1*. Additionally, Tajima’s *D* statistic suggests that *SUL1* is unlikely to be evolving neutrally (*D* = −2.85). This may indicate that this locus has not reached equilibrium after a bottleneck in the past and is still undergoing expansion. The neutrality index calculated from the McDonald-Kreitman test indicated no evidence of selection (NI = 1.117, Fisher’s exact two-tailed test, p = 0.625); however, there are mutations in the *S. cerevisiae* population that are slightly and fully deleterious, which have been shown to cause errors in predictions of adaptive evolution using this test (Charlesworth and Eyre-Walker, 2008).

In order to determine whether *SUL1* is exceptional in the prevalence of loss of function mutations, we determined the frequency of likely deleterious premature stop codons at all loci in the 1,011 strain collection sequences. Using the sequencing data curated in the 1,011 *S. cerevisiae* strains, we analyzed the coding sequences of genes in the pangenome for premature stop codons that occurred in the first 90% of the gene. We excluded genes that either did not appear in the pan-genome or contained premature stop codons in the pan-genome reference sequences. Grouping these genes enriched in premature stop codons by ecological origins, we found that dairy strains tended to have a consistently higher number of genes that are homozygous for premature stop codons compared to strains isolated from other ecological origins (**Supplemental Figure 7**). This is consistent with previous studies that identified enriched loss-of-function alleles among dairy strains that were a result of drift and are important for trait variation (Legras et al., 2018; Zorgo et al., 2012). Of all the genes in the pangenome, 2,465 genes contain a premature stop codon in at least two strains, with 862 of these genes containing premature stop codons in more than 20 strains. Gene Ontology (GO) term analysis revealed that 158 of these 862 genes are involved in ion and/or transmembrane transport. This corresponds with previous analyses that found that genes encoding transmembrane proteins tended to be closer to telomeric ends of chromosomes and were more likely to acquire loss-of-function mutations (Bergström et al., 2014). Of the 1601 genes that have premature stop codons in fewer than 20 strains, 284 are involved in catabolic processes (Holm-Bonferroni test/Benjamini Hochberg p-value < 3e-4) and 385 are involved in responses to stimuli (p-value < 6e-5). The number of genes with loss-of-function variants is much greater than the number found in previous studies, likely due to the fact that this dataset has a greater number of strains and much more diversity among strains in regards to factors such as ploidy and isolation origin (Bergström et al., 2014; Jelier et al., 2011).

## Discussion

Assessing the phenotype of alleles on a species-wide scale is crucial for understanding how quantitative traits vary in a population. Previously developed approaches for experimentally identifying causal variants are conducted through DNA synthesis or mutagenesis, and in many cases do not reflect alleles found in natural populations. We have developed here a high-throughput and low-cost functional approach that can measure the fitness of nearly all alleles present in a population. Specifically in our study, we investigated the function of 228 natural variants of *SUL1*, a high-affinity sulfate transporter gene, present in the 1,011 *S. cerevisiae* strain collection. Our assay identified instances of functional, intermediate, and loss-of-function phenotypes. Using this data, as well as gene and whole genome sequencing data, we related *SUL1* fitness to its evolutionary history. *SUL1* acquired multiple independent instances of loss of function, the majority of which were due to premature stop codons. Other alleles had frameshift, nonsynonymous, and promoter polymorphisms that negatively affected fitness. These multiple independent instances provide evidence that there may be a fitness trade-off for having a loss-of-function *SUL1* allele. The strains carrying these loss-of-function alleles were largely isolated from dairy, beer, and sake clades. Because not all loss-of-function polymorphisms were identical in each clade (for instance, there are three different premature stop codons among dairy strains), these events were likely not due to drift but may have a functional benefit instead. We recognize an alternative possible explanation, which is that some strains, including those from dairy environments, have been shown to naturally carry a high burden of loss of function polymorphisms, and *SUL1* could simply represent an easily tolerated loss that is recurrent by chance. As shown by previous studies, enriched loss-of-function events in specific populations are thought to arise as a result of genetic drift and play an important role in maintaining genetic variation (Legras et al., 2018; Zorgo et al., 2012).

However, there is some evidence that a loss-of-function *SUL1* allele may confer a trade-off and be beneficial under particular environments. Prior studies have shown that there are toxic analogues of sulfate, such as chromate and selenate, that could be transported into the cell through the Sul1 permease (Cherest et al., 1997; Johnson et al., 2016). Several studies have also identified other toxic compounds such as cadmium that affect cell function and growth due to the uptake of sulfate by Sul1 (Zhang et al., 2020). These show instances where having a functional copy of *SUL1* would be detrimental and suggest that *SUL1* may have some antagonistic pleiotropic effects. This may also explain the lack of gain-of-function alleles in our library, as having a higher-affinity *SUL1* may not be beneficial in natural environments. Despite the results from previous studies, many of which investigated the effects of toxic compounds in lab strain backgrounds similar to what we used here, we have been unable to recapitulate these trade-offs.

Identifying loss-of-function alleles by searching for premature stop codons is relatively straightforward. Additionally, we found that many of the nonsynonymous polymorphisms were predicted from mutfunc to have a deleterious effect, although many of these predicted deleterious polymorphisms were false positives. Moreover, the effects of polymorphisms in regulatory regions are more challenging to predict computationally. Using natural variation, we have identified instances where a single polymorphism (n.-456G>A) in a predicted transcription factor-binding site affects fitness of cells under sulfate limitation, a result that was also apparent in our prior promoter mutagenesis study (Rich et al., 2016).

Our approach also identifies intermediate phenotypes, many of which in our pool were likely a result of a natural promoter polymorphism that affects expression. For studying variants, it is challenging to identify deleterious mutations in a population, and here we illustrate an example showing the importance of studying both coding and noncoding polymorphisms, as both normal expression and protein structure affect phenotype and thus how selection acts on a population.

While some *SUL1* alleles have single polymorphisms that can result in a total loss of function, there were also alleles with several nonsynonymous mutations that had wild-type-like fitness under sulfate limitation. Notable examples include the two *SUL1* alleles found across 21 unique isolates that had signatures of introgression from *S. paradoxus*; these alleles had over 40 amino acid differences, yet functioned normally in the S288C background. These results support our previous findings that *SUL1’*s high affinity has been maintained across *S. paradoxus* and *S. cerevisiae* (Sanchez et al., 2017), and the fitness measurements of the introgressed alleles support the idea that these sequences maintain their function even in a new genetic background context. The wide variation in *SUL1* function under sulfate limitation is stark, and using these natural variants has provided further evidence for non-neutral evolution.

In this study and our prior study, we found no correlation between *SUL1* function and its original isolate’s growth on sulfate-limited media (Payen et al., in preparation). Again, despite the fact that *SUL1* copy number increases in evolution experiments under sulfate limitation, we were surprised to see that fitness of endogenous copies of *SUL1* did not necessarily dictate cell performance under sulfate limitation. One possible reason for this observation is that these strains contain functional copies of the *SUL1* paralog, *SUL2*. Despite being a lower functioning sulfate permease compared to *SUL1*, we found no strains that were homozygous for obvious loss-of-function *SUL2* alleles. The alleles of *SUL2* and other transporters like *SOA1* likely also play an important role in growth under sulfate-limiting conditions. Alternatively, small growth rate changes may not be observable in our solid media growth rate assays compared to what is possible to measure in chemostat culture.

All in all, leveraging the technologies available in high-throughput Illumina and PacBio sequencing, we present here a widely applicable and affordable approach for assaying hundreds of natural variants in high-throughput. Assaying natural variants in this manner is especially useful when coupled with whole-genome sequencing data, as it allows us to better understand function in relation to molecular evolution. Furthermore, our method compares many alleles of a gene in isolation in an otherwise isogenic background away from the complexities of genetic background interactions. This approach complements methods like QTL mapping, providing a more thorough investigation of phenotypic patterns across an entire species, which can also contribute to our understanding of how pleiotropic a gene is. Further application of this approach in other genes and other genetic backgrounds will be greatly beneficial to our understanding of how selection acts on natural populations and how multiple polymorphisms contribute to function and ultimately phenotype.

## Acknowledgements

The authors thank members of the Dunham lab for their help: M. Bryce Taylor and Abigail Keller for assisting in experimental design and analysis; Clara Amorosi and Christopher Ryan Livingston Large for helping with quality control and providing input on data visualization; and Noah A. Hanson for sequencing samples. We also thank Anne Clark and Josh Akey for helping with the sequence analysis and identification of introgressed alleles. We are grateful for Mary Kuhner for her consultation and expertise on the best approach for creating and interpreting our phylogenetic trees, and for Pengyao Jiang and Kelley Harris for assisting with visualization of these trees. The authors also thank Fangfei Li and Sasha Levy for patiently assisting us with FitSeq in MATLAB. Additionally, we would like to thank the UW PacBio Sequencing Core for sequencing our samples while also helping us with optimizing the quality of our libraries. This work was supported by the National Science Foundation Graduate Research Fellowship (DGE-1762114). Research reported in this publication was supported by the National Institute of General Medical Sciences of the National Institutes of Health under award number R01GM101091. The research of MJD was supported in part by a Faculty Scholar grant from the Howard Hughes Medical Institute.

---

**Supplemental Figure 1.**
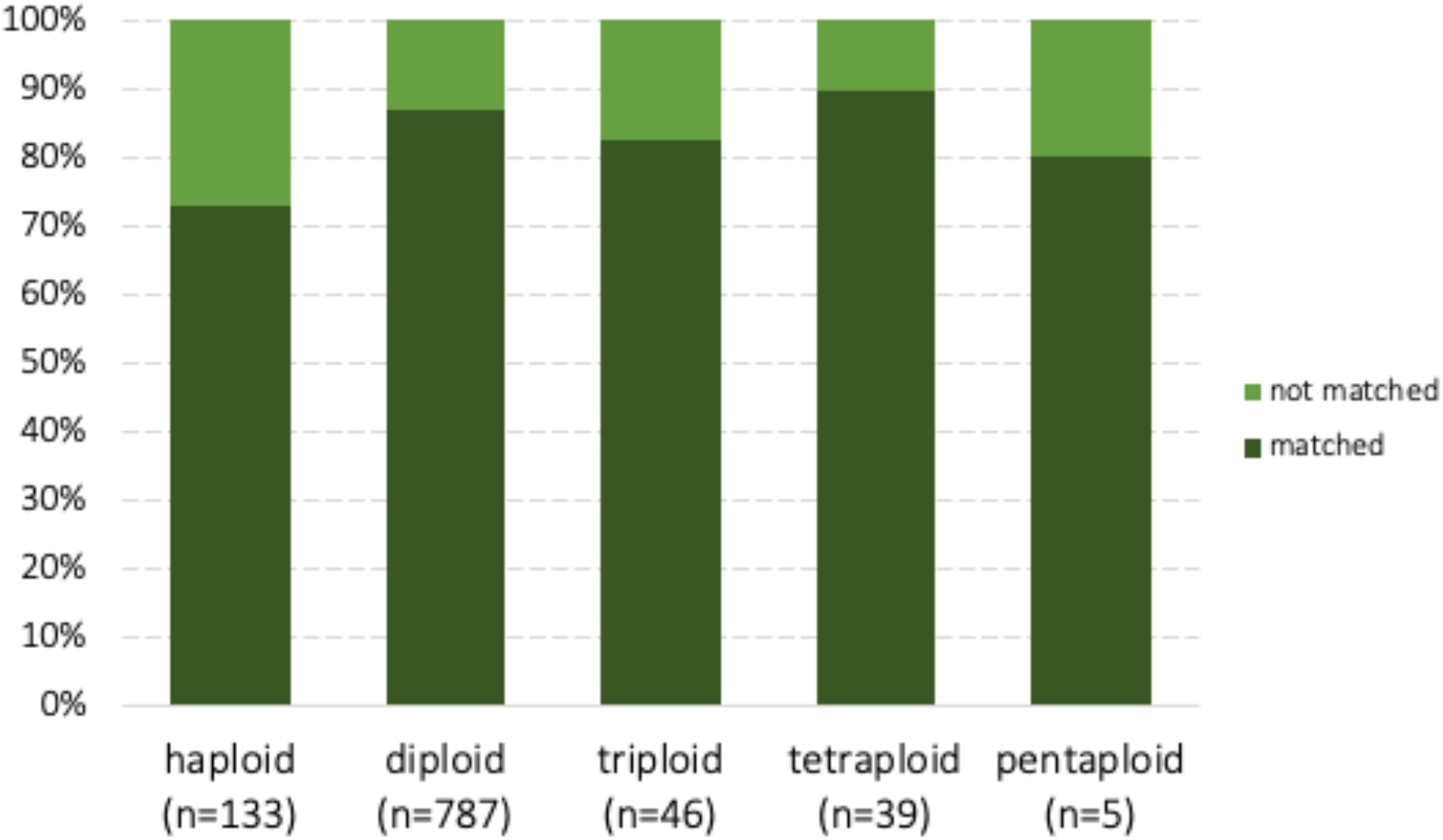
Percentage of strains for each ploidy that matched to at least one PacBio read.

**Supplemental Figure 2.**
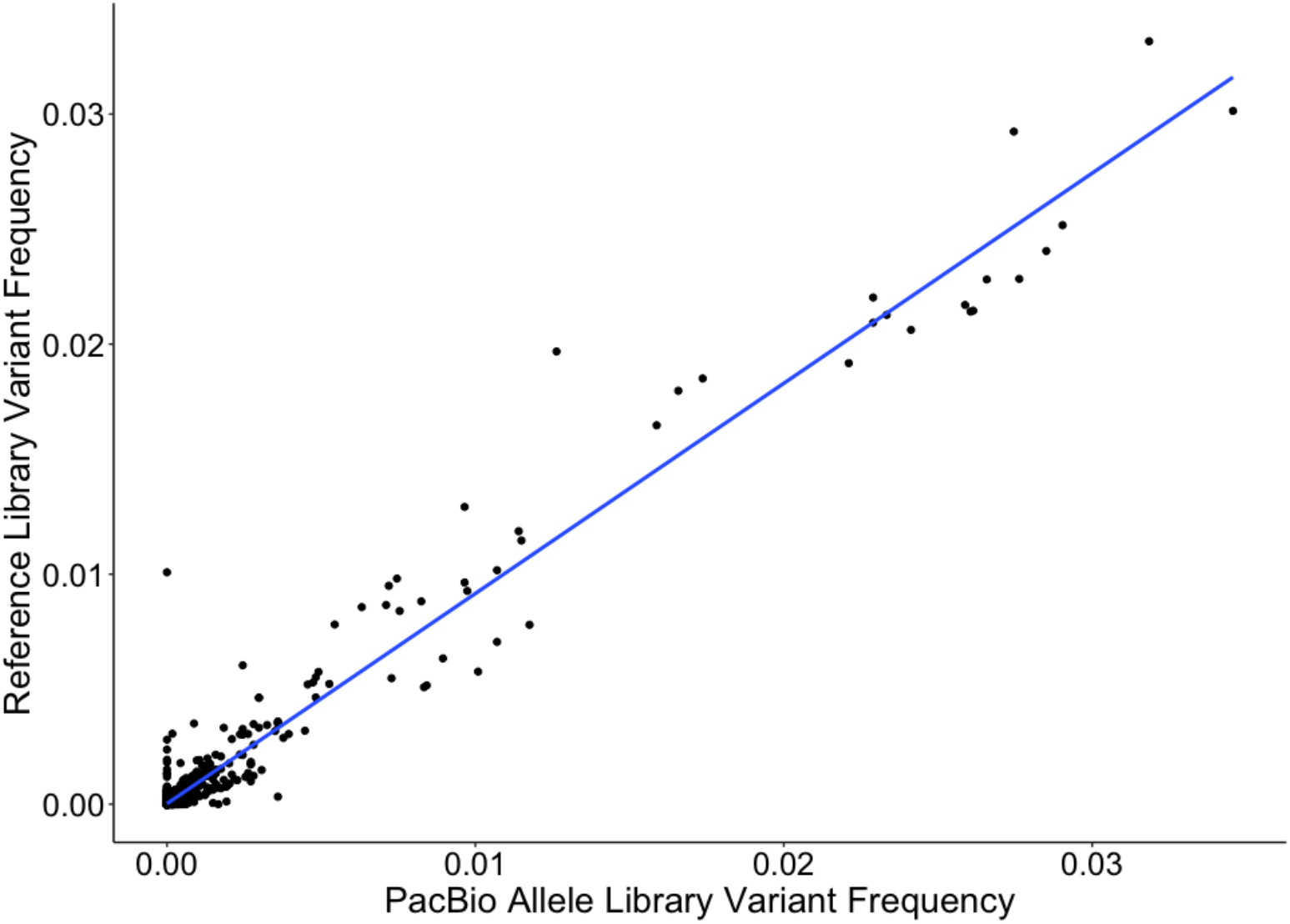
Allele frequencies found in PacBio allele library reflect those found in the Illumina reference sequences (expected values).

**Supplemental Figure 3.**
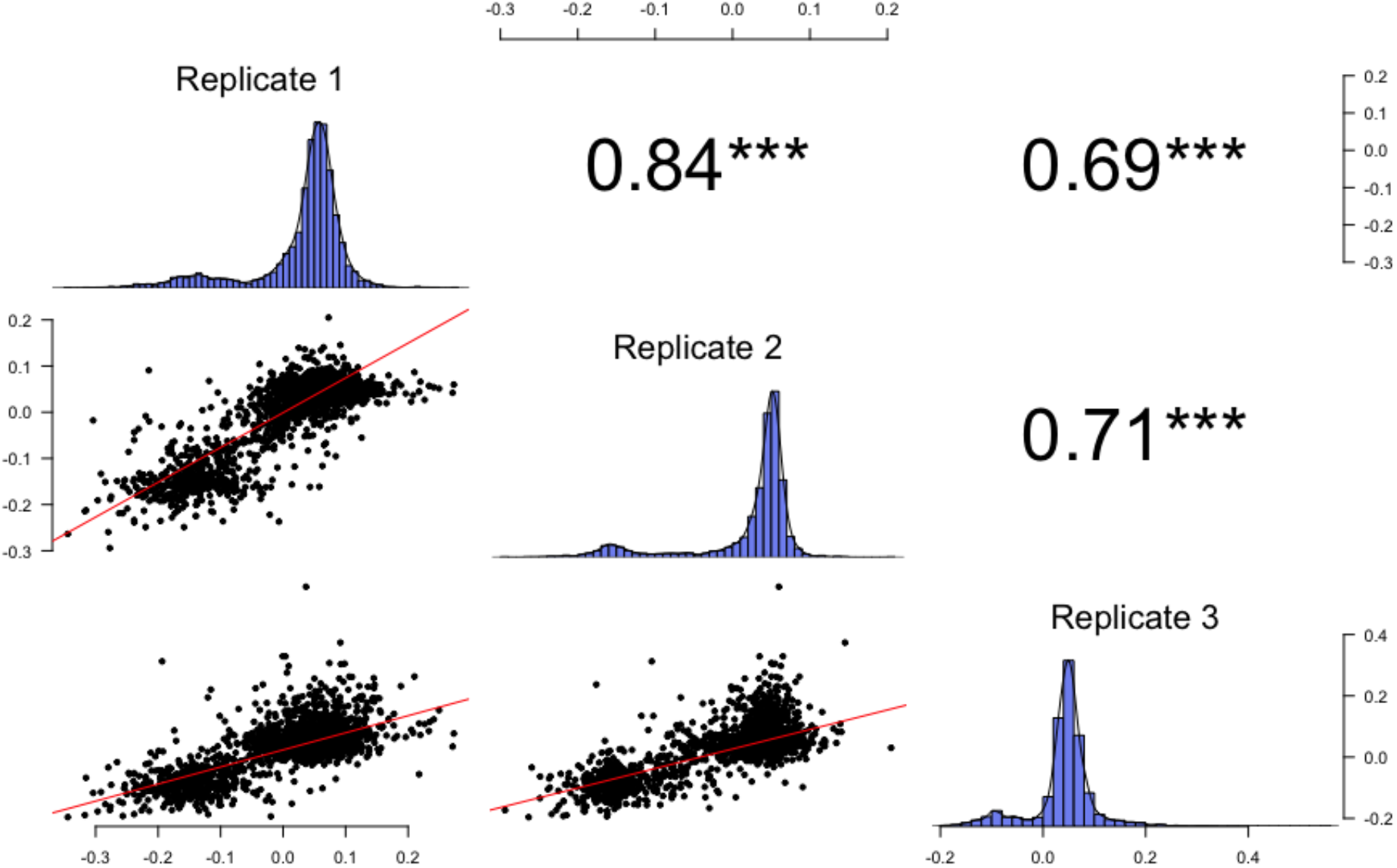
Competitive fitness values calculated using FitSeq are well-correlated across replicates. Pearson correlation coefficients *r* are listed on the top half. ***p<2.2e-16

**Supplemental Figure 4.**
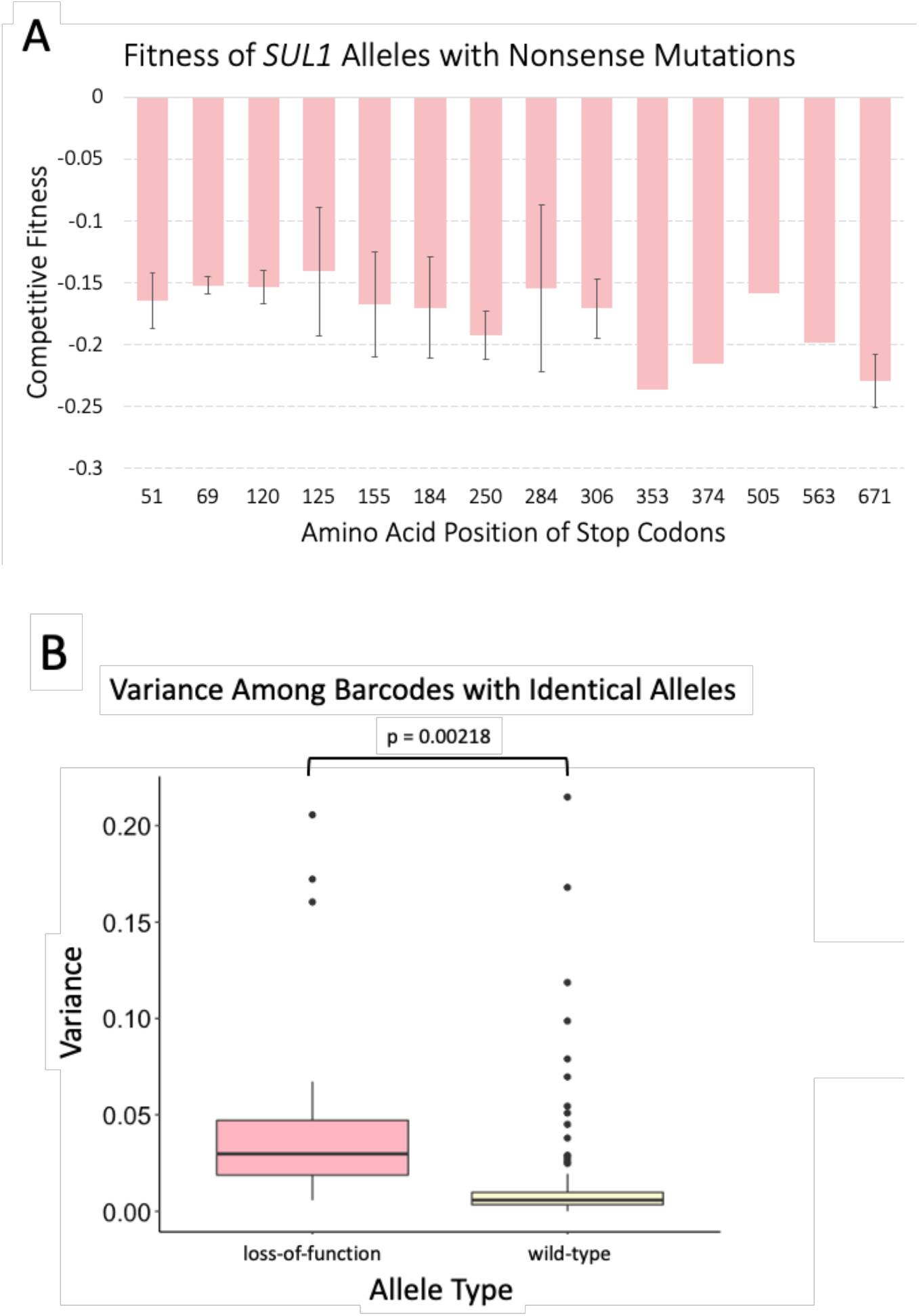
**a)** Barplot representing average fitness and standard deviation of barcodes categorized by location of premature stop codons. Sites without error bars are represented by only one barcode. **b)** Barcodes associated with loss-of-function alleles tend to have greater variance compared to barcodes with wild-type fitness.

**Supplemental Figure 5.**
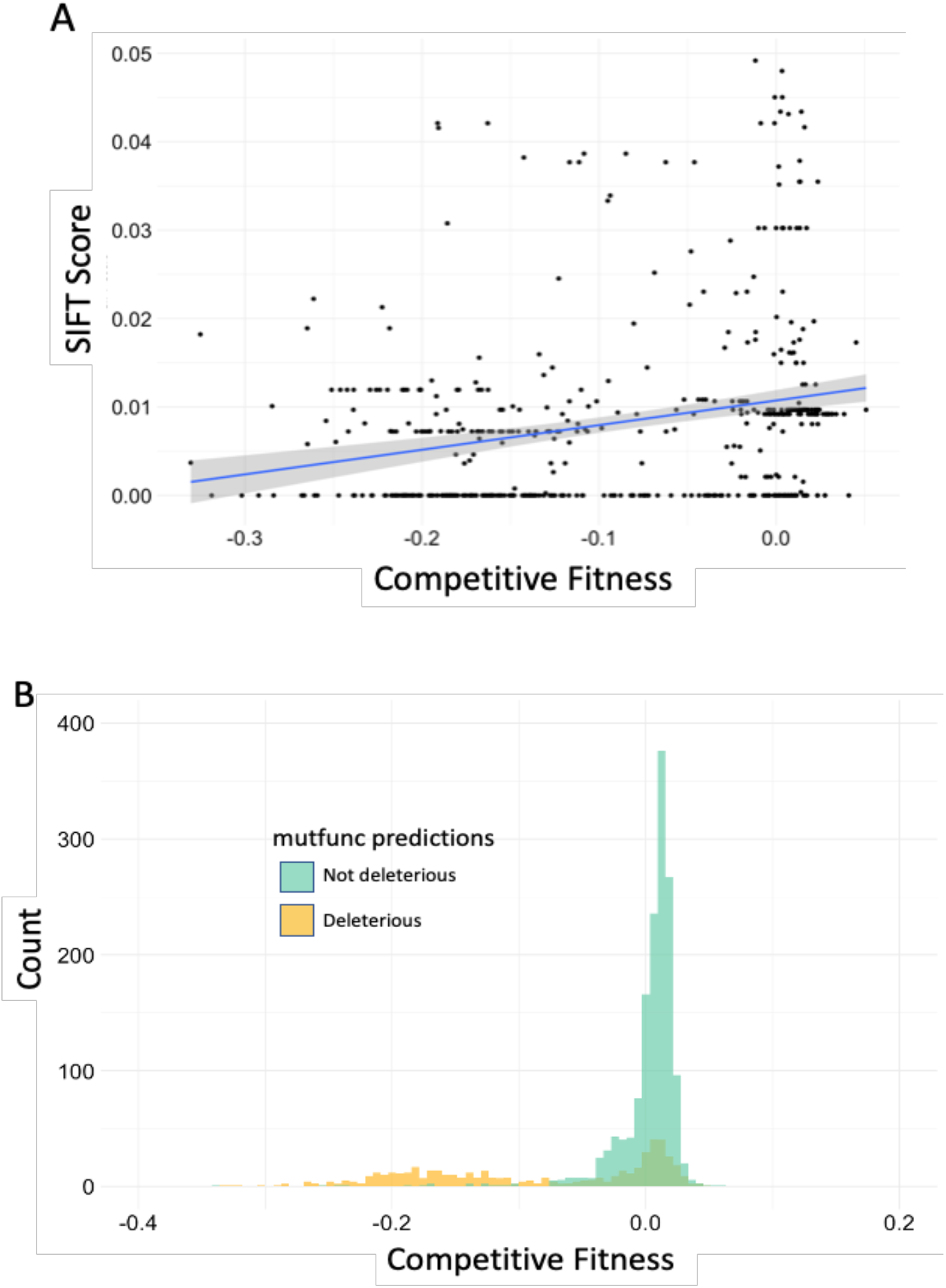
mutfunc determines which mutations are deleterious. For our data, mutfunc returned SIFT scores for each mutation. We used the mutation with the most deleterious SIFT scores for each allele. **a)** Competitive fitness of allele from pooled natural variant library plotted against SIFT score of most deleterious mutation shows very little correlation (Pearson’s correlation r=0.253). **b)** Distribution of experimentally assayed compared with mutfunc predictions of deleteriousness.

**Supplemental Figure 6.**
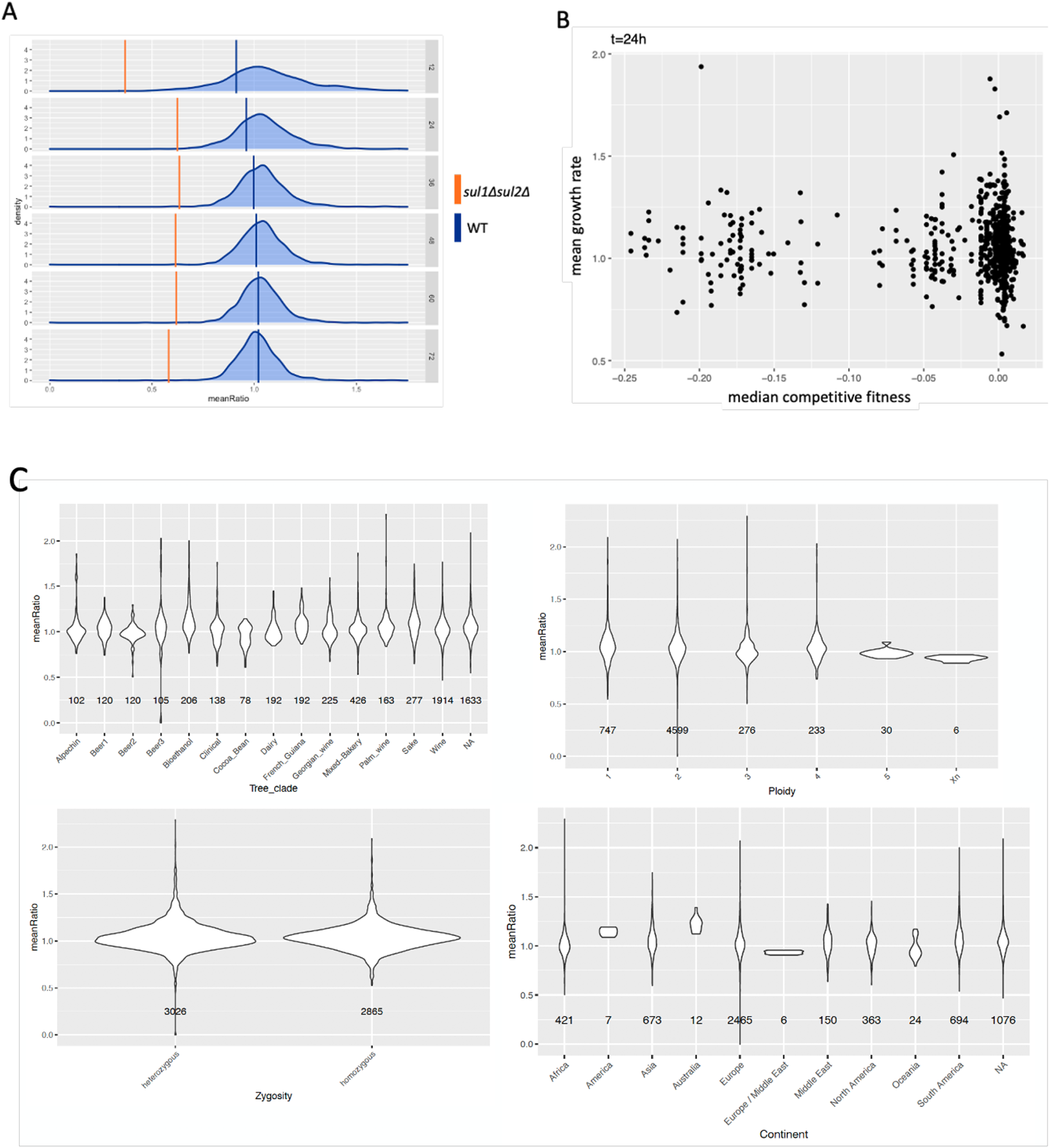
**a)** Growth rate of *sul1Δsul2Δ* strains (orange) and wild-type strain (blue) show differential growth on sulfate-limited media. **b)** Scatterplot comparing strain competitive fitness with growth rate on solid sulfate-limited media show no correlation. **c)** Grouped by clade, ploidy, zygosity, and continent, strains show no obvious pattern

**Supplemental Figure 7.**
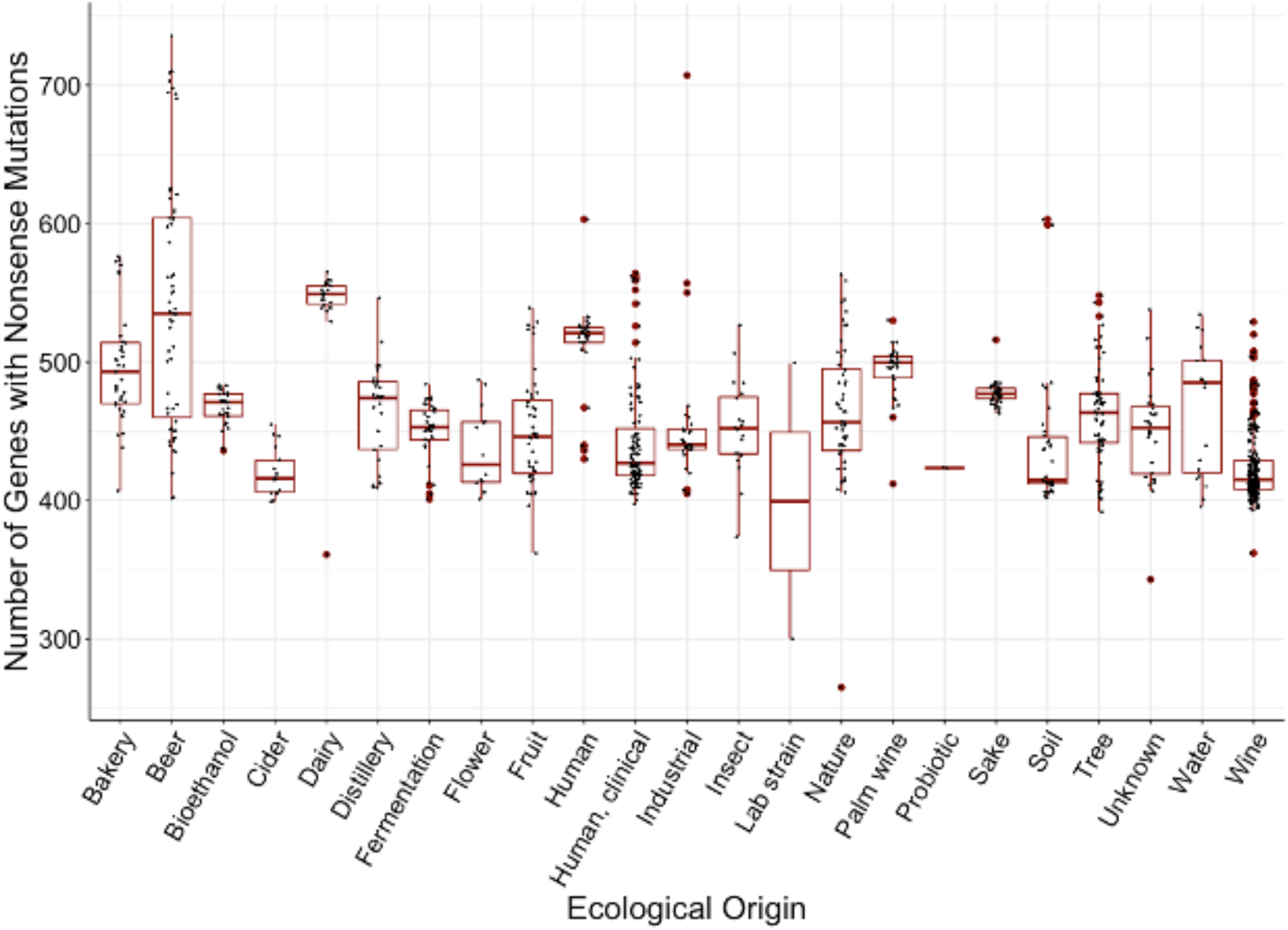
Barplot of number of genes with premature stop codons per strain, grouped by ecological origins.

